# Lessons learned during the journey of data: from experiment to model for predicting kinase affinity, selectivity, polypharmacology, and resistance

**DOI:** 10.1101/2024.09.10.612176

**Authors:** Raquel López-Ríos de Castro, Jaime Rodríguez-Guerra, David Schaller, Talia B. Kimber, Corey Taylor, Jessica B. White, Michael Backenköhler, Alexander Payne, Ben Kaminow, Iván Pulido, Sukrit Singh, Paula Linh Kramer, Guillermo Pérez-Hernández, Andrea Volkamer, John D. Chodera

## Abstract

Recent advances in machine learning (ML) are reshaping drug discovery. Structure-based ML methods use physically-inspired models to predict binding affinities from protein:ligand complexes. These methods promise to enable the integration of data for many related targets, which addresses issues related to data scarcity for single targets and could enable generalizable predictions for a broad range of targets, including mutants. In this work, we report our experiences in building KinoML, a novel framework for ML in target-based small molecule drug discovery with an emphasis on structure-enabled methods. KinoML focuses currently on kinases as the relative structural conservation of this protein superfamily, particularly in the kinase domain, means it is possible to leverage data from the entire superfamily to make structure-informed predictions about binding affinities, selectivities, and drug resistance. Some key lessons learned in building KinoML include: the importance of reproducible data collection and deposition, the harmonization of molecular data and featurization, and the choice of the right data format to ensure reusability and reproducibility of ML models. As a result, KinoML allows users to easily achieve three tasks: accessing and curating molecular data; featurizing this data with representations suitable for ML applications; and running reproducible ML experiments that require access to ligand, protein, and assay information to predict ligand affinity. Despite KinoML focusing on kinases, this framework can be applied to other proteins. The lessons reported here can help guide the development of platforms for structure-enabled ML in other areas of drug discovery.

## Introduction

Phosphorylation, or the transfer of an ATP-derived phosphate group to substrate proteins, lipids, or carbohydrates, is one of the most common post-translational modifications and potentiates a wide variety of intracellular signaling cascades. [1, 2] This reaction is catalyzed by a class of enzymes called kinases, of which there are nearly 540 unique human proteins. [3] Given the centrality of phosphorylation in growth, proliferation, motility, differentiation, and other essential biological processes, kinases are implicated in a variety of diseases and are an important drug target, particularly in oncology indications. Since the approval of the first kinase inhibitor, imatinib, in 2001 through 2023, the U.S. Food and Drug Administration (FDA) has approved more than 80 small molecule protein kinase inhibitors. [4]

The pharmacological importance of kinases has led to their robust characterization and the generation of a vast amount of pharmacokinetic, biochemical, and structural data relating to their function and inhibition. While this makes kinases an ideal protein family to interrogate with machine learning (ML) methods as drug targets, this abundance also poses a challenge to curating a high-quality data set that reflects the breadth and scope of available data while ensuring its accuracy, consistency, reproducibility, and availability.

Both ligand-based and structure-based ML methods are nowadays commonly used in kinase drug discovery. [5–10] Ligand-based methods rely on the “similarity-property principle” [11] stating that ligands with similar structural characteristics will display similar binding affinities to the same target. Therefore, these methods require a set of compounds with known—and ideally varying—activity against a target of interest. Furthermore, using structure-activity relationships (SAR) these compounds can be improved by designing appropriate analogs. [6, 11, 12] However, ligand-based methods do not explicitly include the 3D structural information of the protein:ligand complex. Therefore, they cannot provide direct insights into the structural features of protein:ligand molecular interactions. [8] On the other hand, structure-based methods use protein:ligand 3D structural and interaction data to make binding affinity predictions. [8, 13] It is hypothesized that incorporating structural data into the ML model could improve the binding affinity predictions [9, 13], thus, potentially exceeding the performance of ligand-based methods.

### Why structure-based methods?

Structure-based ML models use the structural information of protein:ligand complexes to predict ligand binding affinities from the ligand’s interactions with the protein binding site, whereas ligand-based models use information derived only from the chemical structure of the ligand. [5, 9, 14]. A key difference between the two approaches is that structure-based methods are able to integrate data from related targets into a single unified model, while ligand-based methods generally do not transfer well to related targets. We hypothesize that **by training a structure-based, kinome-wide model with all compound data available for any kinase, the model will learn the physics of protein:ligand interactions well enough to explore larger areas of chemical space than ligand-based methods do**.[5, 14]. Furthermore, although there is more available compound affinity data than complex structure data, leveraging all available kinase data for all targets may overcome this limitation.

We note that there is a long history of ML models that incorporate protein information (usually the sequence) without including the 3D structure, which also benefit from incorporating compound data for multiple targets. Two examples are drug-targeting interaction (DTI) and ProteoChemometric modeling (PCM) [15, 16]. However, **by considering the precise interactions between the ligand and the protein binding pocket from the atomic representation, structure-based models may better capture the binding mechanisms**. [5] Including this information in ML models should improve binding affinity prediction compared to ligand-based methods [8, 13]. Taking into account the protein:ligand interactions enables the detection of structural differences that lead to variations in binding free energies, as well as molecular differences between binding modes of ligands across a range of kinases. We hypothesize that this will lead to improved performance in optimizing ligand potency and predicting selectivity, mutational resistance, and polypharmacology. For example, *Backenkohler et al*. [9] showed improved performance in binding affinity prediction for kinase inhibitors of their structure-based model, over 3D-structure-free models such as DTI. Similarly, *Luo et al*. also showed improved performance in kinase-drug binding affinity prediction of their structure-based method over structure-free methods [17].

Finally, like many proteins, kinases can exist in multiple conformational states, which can be clustere into e.g., the four KLIFS states [18] or the eight Dunbrack states [19, 20]. **The conformational dynamics of kinases may be a key to the design of effective kinase drugs, and structure-based models have the potential to encode this conformational information and use it to predict binding affinities**. This again would be a dimension missing from ligand-based methods.

### Overcoming the data challenges

ML, particularly deep learning (DL), models are very data greedy since they derive patterns solely based on the given training data. In order to obtain good performance a large amount of data is needed. [15] In the case of kinases, there are various databases available that collect kinase information including structural protein:ligand and specific annotation data. KLIFS [21], e.g., hosts over 6, 667 PDBs of kinases and unique kinase:ligand combinations [18]. Particularly, there are currently 4, 148 unique kinase inhibitors in KLIFS. While this amount of structural data is large for a protein family, it is not enough for reliable DL applications. Therefore, structure-based ML models in the field of inhibitor design face the challenge of limited available structural data of protein:ligand complexes with experimental affinity data, especially compared to the relatively more available data for ligand-based methods.

In order to exploit the full potential of ML structure-based models, structural protein-ligand data for unresolved structures needs to be generated. To tackle this, there have been several initiatives focused on data augmentation and structure predictions. For example, the KinCo [8] data set comprises a kinase-inhibitor complex data set of 137, 778 *in silico* predicted kinase:ligand structures paired with experimental binding constants. The kinase structures were predicted using homology modelling, and then compounds were docked to these structures. Furthermore, Schaller et al. [13] introduced a kinase-centric template docking workflow to augment the structurally accessible kinase-ligand space with promising performance in a benchmark study. More recently, Backenköhler et al. [9] built on this study, and used the template docking workflow to generate structural information of all kinase:ligand pairs with available bioactivity data in ChEMBL. Overall, approximately 140, 977 kinase:ligand pairs were obtained and evaluated. Notably, the study showed a significant increase in model performance, when trained on this dataset and using the complex 3D information encoded in an E(3)-invariant GNN model.

In the following sections, we report our experiences in building KinoML. KinoML is a modular and extensible framework for machine learning (ML) in small molecule drug discovery with a special focus on kinases. The purpose of KinoML is to help users conduct ML kinase experiments, from data collection to model evaluation. Note that despite KinoML’s focus being on kinases, it can be applied to any protein system. Tutorials on how to use KinoML as well as working examples showcasing how to use KinoML to perform experiments end-to-end can be found here.

In order to describe our experiences in building such a framework, we will first dive into how to obtain and deposit data from different sources in a FAIR and reproducible way, including how to preprocess the data properly. Secondly, we will introduce KinoML, with a description of its main goals and object model along with example scripts explaining how to use it. This will be followed by a description of how to store the information as tensors for easy ingestion in ML applications. To conclude, we summarize the key learnings to conduct ML-based kinase drug discovery experiments transferable also to other areas of drug discovery.

## The data journey to KinoData

In this section, we will discuss how to collect, curate, integrate, and harmonize kinase data from different sources in a reproducible way and with an emphasis on FAIR principles. These efforts resulted in **KinoData**, which is a collection of Jupyter notebooks to reliably select and curate datasets. These scripts can be found here.

### Identifying and integrating useful data sources in a reproducible way

High-throughput kinase-ligand binding affinity experiments are routinely conducted, generating valuable data for individual research questions but also for ML models and large-scale simulations. For instance, standardized assay platforms like DiscoverX^1^, KinomeSCAN^2^, and KdELECT^3^ provide experimental binding affinity values for kinase:ligand pairs. Also, regarding only kinases, there is publicly available data such as the Published Kinase Inhibitor Set 2 (PKIS2). [22] Furthermore, there are publicly available curated databases of bioactivity of molecules with drug-like properties, that are not specific to kinases, such as ChEMBL^4^[23] or PubChem^5^.

However, proper data handling (data preprocessing) is crucial before integrating data into any pipeline. This involves deduplication, unit system standardization, mislabeled entry filtering, and other data-wrangling tasks that significantly impact downstream model and simulation quality. Once the data artifacts have been identified and removed, they need to be archived for provenance and reproducibility. In this section, we will discuss key considerations for building a reproducible data generation process from existing biomedical resources. For this purpose, this section will cover good practices on how to collect data and also on how to deposit this data. With this in mind, we also introduce here our KinoData project ^6^, which consists of a series of scripts that exemplify how to reliably select and curate kinase-related data sets.

#### Reproducible data collection

Our journey starts with obtaining data from the chosen data source(s). It is a common misconception that data acquired at the project’s outset will remain available and unaltered throughout the experiment’s lifespan. Many data repositories will undergo updates and changes, making it necessary to take proactive steps to ensure our data sources exhibit two properties essential for reproducibility: *availability* and *immutability*. These characteristics are vital to ensure that we or others can reproduce our findings in the future. There are three main sources to obtain raw molecular data: online data sets, peer-reviewed publications (with supporting information), and shared files.

##### Online data sets

Public repositories of biochemical data like ChEMBL^7^ and PubChem^8^ provide frequently updated data accessible on demand. Therefore, automated retrieval through APIs does not guarantee identical results over time due to continuous changes in these databases. To address this, we recommend using versioned database copies, which are often available through official download portals in various formats (e.g., SQLite, MySQL). These archived versions, accessible via Digital Object Identifier (DOI) links, exemplify immutable availability. If your data source lacks versioned downloads, consider creating your own local copy via APIs or web scraping, and depositing it in online archives like FigShare, Zenodo, or GitHub Releases if permissible. An example of querying human kinase inhibitor bioactivity data from a specific ChEMBL version can be found in the KinoData tutorial *kinases_in_chembl*.*ipynb* ^9^.

##### Peer-reviewed publications or shared files

Some data sets are deposited as supporting information in peer-reviewed publications with DOIs, such as PKIS2. [22] Availability is ensured through the publisher, but differences in how files are provided may make automated retrieval challenging. If specific data is not publicly accessible, it may be necessary to request it from another researcher (if there is a Data Availability Statement). Alternatively, if files are shared through private channels like email or Slack, ensure their public deposition with a permanent URL to achieve long-term availability and immutability. Another point to keep in mind is that licensing issues may be a hindrance when using data from other researchers or organizations. In this case, it is recommended to directly contact the authors. Moreover, when considering which licensing scheme to use for your own data, it is recommended to use the CC0 (public domain) license [24]. This type of licensing allows data to be treated as if it is in the public domain, facilitating aggregation of data into other repositories, thereby promoting collaborations and knowledge sharing.

Following these recommendations when dealing with online data sets, peer-reviewed publications, or privately shared files will ensure that the data remains available and unchanged, promoting the reproducibility of your research.

#### Ensuring your processed data is FAIR

The FAIR principles are a set of guidelines for researchers to ensure their data is **F**indable, **A**ccessible, **I**nteroperable, and **R**eusable [25] and are key for open science. Following these recommendations allows scientists to reliably access data, combine data from different sources, and ease reproducibility of results. In this section, we cover the key steps to ensure compliance with FAIR guidelines once the data has been collected.

Normally, raw data should be preprocessed before archiving it to save resources. This is particularly important when dealing with large data sets or when data is obtained from non-API sources. If data preprocessing is necessary (e.g., extracting data from HTML), it is essential to include the corresponding code alongside the final data set. This ensures that updating the archived data only requires rerunning the preprocessing pipeline and enables the community to understand any preprocessing applied to the data. Three key qualities for deposited data are a unique identifier, guaranteed long-term availability, and accessibility for both human and programmatic interfaces.

The choice of where to store your data set depends on its size. For *small data sets* (a few megabytes), standard version control systems like GitHub can be sufficient. These systems provide unique identifiers that can be accessed via URLs, typically using commit hashes or tags. For *medium-sized data sets* (several hundred megabytes), you can use platforms like GitHub Releases. This approach involves publishing a tagged commit as a “release” and allows you to attach the data set as a separate artifact. It is advisable to upload the data set in a compressed format, which can be done manually or automatically through Continuous Integration (CI) pipelines like GitHub Actions. However, it is important to always keep in mind that GitHub URLs are not guaranteed to be permanent; for example, they can disappear due to the repository being deleted by the owner. Therefore, when sharing and storing GitHub URLs, it is recommended to use platforms like Zenodo^10^ (for versioned releases and DOI assignment). Finally, for *very large data sets*, often exceeding the file size limits of version control systems (in the order of a few gigabytes), external cloud storage providers like FigShare^11^ or Zenodo are recommended. These platforms can handle large data sets and also support DOIs if needed.

### Reproducible data collection and deposition example: KinoData

In this section, we illustrate our data processing pipeline—data set identification, ingestion, and deposition— with kinase inhibitor activity data information in a modular and reproducible way. This is divided into three steps: first, identifying the relevant target kinases; second, collecting the compound and bioactivity data; and third, curating the database for its deposition. This process will be illustrated using examples from KinoData.

#### 1. Data identification: what is a kinase?

The first step is to retrieve the kinase data of interest from the different databases. However, the definition of *kinase* is not as straightforward as one would expect. Throughout the literature, one can find several “authoritative” lists of kinases. Unfortunately, the criteria used to nominate how a protein makes it to the kinase list is neither consistent nor obvious, resulting in divergent definitions of what the human kinome really represents. Of course, this also hinders the querying of kinase-related data from data sources. To address this issue, we referred to several publications and the most relevant kinase-centric web-servers. The kinase-centric data sources studied were:

- KinHub is a web-based tool for interactive navigation through and visualization of human kinome data, following the nomenclature by Manning et al. [26]. Pre-processed data from freely available sources such as ChEMBL [23], the Protein Data Bank (PDB) (https://www.rcsb.org/), and the Center for Therapeutic Target Validation (CTTV) were compiled, integrated, and visualized on the kinome tree. [27]
- KLIFS [18]: The Kinase–Ligand Interaction Fingerprints and Structures (KLIFS) database collects all kinase structures and aligns, annotates, and curates them with related kinase information. KLIFS now contains over 6, 667 annotated structures, comprised of 326 unique kinases and over 4, 148 unique ligands as of May 2024.
- KinCoRe [28] is a web resource that collects and curates all protein kinase structures from the PDB (https://www.rcsb.org/). KinCoRe assigns conformational and inhibitor type labels that reflect the diversity of the kinase conformations beyond DFG-in/out and inhibitor binding locations.

Several nomenclatures exist to refer to kinases, i.e., xName (KinHub), Manning’s kinase classification [29], or HUGO Gene Nomenclature Committee (HGNC) [30]. While there are decent overlaps in the names, there is no uniquely identifying kinase naming system. Therefore, it is advisable to use Uniprot identifiers as the primary search keys when combining and comparing proteins across different data sources. A Uniprot “Entry name” is a unique and universal alphanumerical code assigned to a specific protein [31, 32]. The resulting set of human kinases, which arises from combining the different kinase definitions from the sources mentioned above, with their different nomenclature and database keys are all available in KinoData in a Jupyter Notebook stored here for permanent availability. The notebook illustrates how to obtain this multisource kinase data step by step, depending on several online resources. The search and mapping resulted in around 500 kinases per set, of which 473 appeared in all five queried data sets (see Figure 1.a).

**Figure 1.**
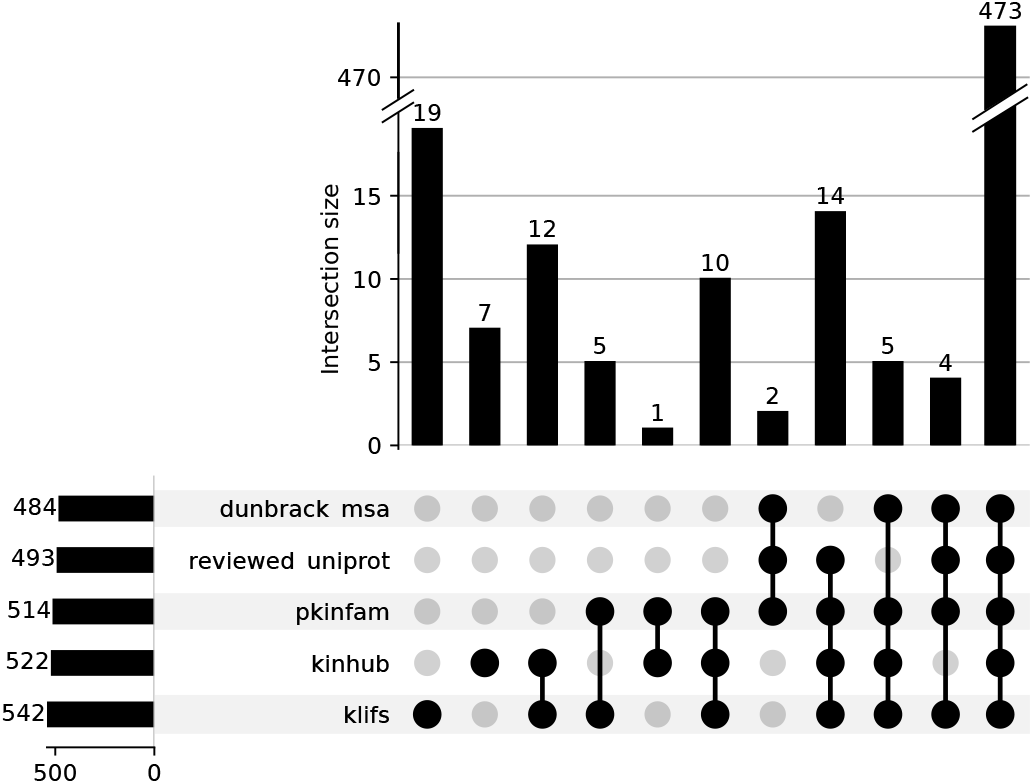
Concordance of definitions of human kinases from various sources reporting members of the human kinome. Data assembled and figure created in https://github.com/openkinome/kinodata/blob/master/human-kinases/human_kinases.ipynb. Overlap of human kinases queried from different sources.

#### 2. Data collection and curation: bioactivity data

Once the kinase target set is obtained, the next step is to obtain bioactivity measurements for these kinases from online databases or other resources. For this purpose, we considered two online databases: ChEMBL [23] and PKIS2 [22]. In this section, we discuss the advantages and disadvantages of ChEMBL and PKIS2, followed by a step-by-step example of how to collect and curate ChEMBL kinase data using KinoML.

##### ChEMBL

ChEMBL [23] is one of the largest and most well-known databases containing bioactivity data for over 2.4 million distinct compounds (as of May 2024) and is therefore of interest for any kinase-centric ML study. Given a kinase target set, querying bioactivity measurements in ChEMBL can be done with the UniProt ID, since ChEMBL offers a mapping from UniProt identifiers to their own internal labels (ChEMBL target IDs). In ChEMBL, a target can be part of different kinds of assays, and thus different types of measurements are reported. For the purpose of this study, we are only interested in single protein measurements, i.e., binding assays with *K*__*d*__, *IC*50, or *K*__;__ data available.

Due to technical limitations in the ChEMBL API, which rate limits big queries, we settled for querying a local copy of the database, as available through DOI http://doi.org/10.6019/CHEMBL.database.33. As mentioned earlier, a local copy provides better performance and reproducibility, since it is immutable and versioned. An example of how to filter and process kinase data from ChEMBL can be found in the following Jupyter Notebook from KinoData ^12^. In this specific example, the resulting data set was loaded in a Pandas dataframe which was filtered following previously published advice [33, 34]. Figure 2 shows the number of PDBs at each step of the ChEMBL curation. Firstly, the data was grouped by protein and ligand, which resulted in 273, 555 ChEMBL measurements. We call a protein and ligand complex a system. Also, extreme affinity values (> 10mM or < 1fM) or measurements with unclear units were removed, resulting in 272, 265 ChEMBL measurements, as depicted in Figure 2. For systems with several measurements in the same publication, only the highest measured affinity was kept. Two final stages removed duplicates where a manuscript cites the original experimental publication and also in cases of author overlap where the same system was referenced in multiple publications. All these steps resulted in a final 211, 697 measurements as seen in Figure 2.

**Figure 2.**
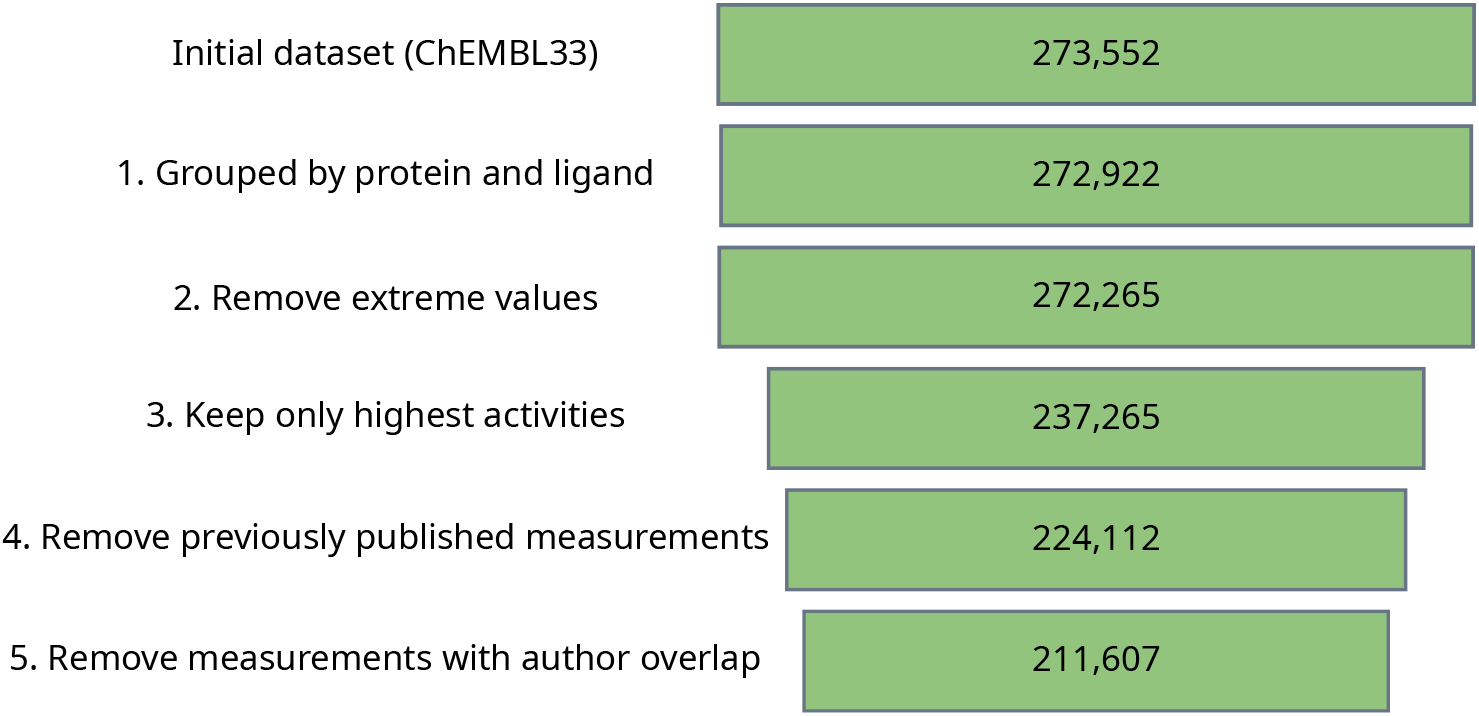
Applying an automated curation pipeline reduces error and safeguards its reliability. 1. Measurements were grouped by their protein (ChEMBL target) and ligand (compound identifiers). A ‘dummy’ ChEMBL target (CHEMBL612545), used to verify unchecked analyses; is removed. 2. All activities were converted from K_i_ to pK_i_ and extremevalues (> 10mM or < 1fM) were removed; 3. Where a target or compound had multiple, different activity values inthe same publication, only the highest pK_i_ was retained. This step was necessary to remove unclear stereoisomer annotationsand measurements taken as part of assay optimizations; 4. Citations within a publication of previously published (i.e. identical) values were removed; 5. Measurements for the same system from different publications if they share one or more authors were removed to identify truly independent pairs of measurements.

##### PKIS2

Another very popular kinase data set considered was Published Kinase Inhibitor Set 2 (PKIS2) [22], which is a different type of data set. It is a published and static document, distributed as a spreadsheet, that does not change with time. It provides a single type of measurement (% displacement) for the inhibitory power of a library of compounds against a panel of human kinases. The compounds are listed as simplified molecular-input line-entry system (SMILES) and the kinases are identified by *name*.

However, there are several challenges and questions behind that simple scheme, such as the name being used in each assay. For example, it was not clear if it was the gene name or the protein name. To answer this question, we contacted DiscoverX, manufacturer of the KinomeSCAN assay kit used in PKIS2 to elucidate which exact constructs are behind each kinase name. DiscoverX generously provided us with a spreadsheet that contained additional information about each kinase in their kit. Still, the data needed further processing to map the data in PKIS2 to the UniProt identifier of each kinase name. This allowed us to disambiguate the kinase names and provide the identifiers and sequences associated with each data point.

#### 3. Data set deposition

Finally, once all measurements of interest for the target kinases were obtained, the data was deposited in KinoData https://github.com/openkinome/kinodata. This way, KinoData contains all steps needed to collect and deposit kinase-centric data and the data is also version controlled available.

### Harmonizing data representations: Assay classes and molecular entities

Once the data is cleaned, we need to consider multiple sources of heterogeneity in the resulting measurement data. KinoML enables users to address some of these issues directly by retaining identifier, construct, and available assay condition in its data objects. However, other concerns will need to be considered by users when curating data sets and building models. Here we summarize some kinase and inhibitor-specific considerations for users to contemplate.

#### Kinase target considerations

Standardizing the kinase target data requires accounting for disparate naming conventions and, where possible, retaining relevant experimental technical information that may influence assay results.

- **Identifier disambiguation:**The identifiers that map the gene product assayed to the source protein sequence may differ between data sources. For example, ChEMBL disambiguates target proteins using Uniprot IDs. In contrast, the PKIS2’s KinomeScan assay uses National Center for Biotechnology Information (NCBI) Accession Numbers to map to protein sequences but the percentage displacement measurements are classified by DiscoverX gene symbols which closely conform to HGNC gene names. We retain both identifiers in our PKIS2 `DatasetProvider` object, discussed later. Cross-referencing these target identifiers may require querying additional database resources.
- **Construct differences:**The protein constructs are the specific forms of the peptide utilized experimentally may differ both from the canonical endogenous protein and between studies. Mutant constructs or those containing different functional domains may demonstrate distinct small molecules binding affnities. Full-length kinase constructs are not generally synthesized for *in vitro* biochemical assays. Instead, the conserved, catalytically active portion of the protein known as the kinase domain is often generated. [35] Post-translational modifications to the underlying construct, particularly phosphorylation in the case of kinases, can also influence intrinsic activity.
- **Assay conditions:**Assay conditions can also influence a target’s form and function and the resulting assay measurements. For example, the experimental pH and redox conditions can determine whether adjacent cysteine residues form disulfide bridges. [36] Similarly, the presence or absence of other proteins (e.g., cyclins in the case of cyclin-dependent kinases) may modulate construct catalytic activity. Finally, the use of a metal cofactor can also alter binding affinity; endogenously magnesium is the preferred kinase cofactor to catalyze phosphorylation, but other divalent ions like manganese and zinc may also be used experimentally. [37]

#### Ligand considerations

Harmonizing ligand data poses challenges analogous to the kinase targets while along unique concerns, including a generally less standardized identifier generation process and the need to remedy of non-canonical or degenerate molecular representations.

- **Identifier disambiguation:**As with the targets, which ligand identifier is used may vary across data sources. PKIS2 provides compound names generated by suppliers or manufacturers (e.g., Glaxo-SmithKline). For drugs without PKIS-supplied names, SMILES strings are used instead. In contrast, ChEMBL uses its own ChEMBL identifiers. In general, small molecule nomenclature depends on the stage of development to which the drug was advanced. Substances that are strictly tool compounds for preclinical applications may possess only a manufacturer-generated internal identifier. An investigational agent that advances to clinical trials may be granted a standardized, generic international non-proprietary name (INN) by the World Health Organization. [38] Marketed drugs may also possess brand names that can vary by geography.
- **Canonical representation:**Once ligand identity has been ingested, a representation amenable to ML applications must be obtained. SMILES strings are perhaps the most commonly used molecular representation, but they are not unique identifiers. The same molecule can possess multiple valid SMILES string representations. As such, cheminformatics packages like RDKit also implement canonical SMILES string generation to enable consistency when determining which of the possible representations should be used. In both ChEMBL and PKIS2, mapping from identifiers to sequences is simplified by the provision of SMILES strings by each database directly. In the case of molecules with chiral centers, some ligands may require further stereochemical specification or may even exist in multiple distinct configurations in the case of racemic mixtures. Other techniques for representing molecules in data sets include molecular fingerprinting, which encodes chemical information in a 1D vector, or graph structure, where constituent atoms and their corresponding bonds are represented as vertices and edges, respectively. [39]
- **Assay conditions:**Molecules may possess different salt forms, whose solubilities and dissolution rates can impact the real administered dose. Additionally, a ligand’s ionization state depends on the pH of the solution and the pKa of the ligand, and its tautomerization states can be influenced by temperature, solvent, pH, and the presence or absence of catalytic acids or bases. [40]

### Retaining and modeling assay information

#### Assay conditions

As discussed above, a variety of assay conditions can alter the underlying biochemical properties of a ligand and its target and influence their measured affinity. When modeling the outcomes of these assays, these experimental parameters should be treated as covariates within the model. For example, pH influences, among other things, the presence of disulfide bridges among the cysteine residues of the target and potential ligand ionization and tautomerization. All of these factors may significantly impact the measured affinity of the target for the ligand; hence, an otherwise identical system assayed under acidic and basic conditions could produce drastically different results. The pH at which the ChEMBL and PKIS2 assays were conducted (pH 7) are captured for the processed data sets as `AssayConditions` in the KinoML object model, which is discussed in further detail in the next section.

#### Assay class

The assays used to assess binding across different data set may measure distinct quantities. For example, some assays may provide direct readouts of binding affinities (K__d__), indirect biochemical readouts of apparent inhibition constants (K__i__), concentrations representing biochemical inhibition (IC__50__), or other readouts of binding (e.g., single-concentration % displacement for a KinomeSCAN assay). The relative frequency of these measurement types in the processed PKIS2 and ChEMBL data sets is shown in Figure 3. One approach to modeling similar but non-identical biochemical assay measurements would be deterministically converting the outcomes to a single, unified measurement type on which the model is trained.

**Figure 3.**
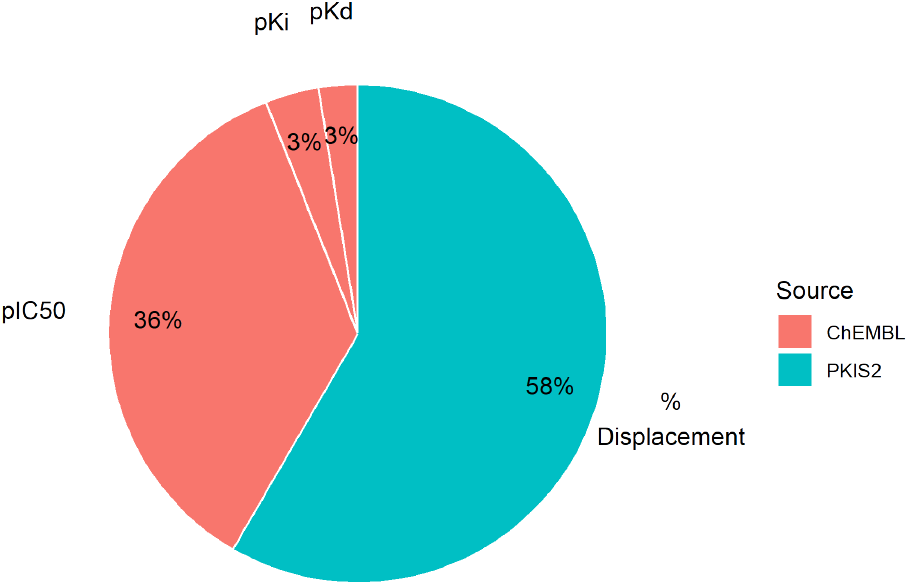
Bioactivity assay measurement classes vary by and within data set. While both ChEMBL and PKIS2 possess kinase bioactivity assay data, the experimental quantity measured differs. The `kinoml.data sets.chembl` object contains 159; 823 IC_50_, 15; 578 K_i_, and 11; 412 K_d_ measurements. The `kinoml.data sets.pkis2` object contains 261; 870 percentage displacement measurements.

Alternatively, rather than transforming the underlying data to a single, standardized measurement type prior to training an ML model, the model’s loss function can be adapted to account for differing measurement classes and conform the output to the input for the purposes of optimization. For example, in the case of a neural network model, the second to last layer could be used to predict a thermodynamic quantity such as Gibbs free energy (Δ*G*) that has a deterministic relationship to the various measurement classes. As long as this conversion function in the final layer is differentiable, the ability of the network to backpropagate and effectively learn parameter weights remains intact.

### From data to numbers: KinoML

To account for the experimental diversity across kinase inhibitor binding affinities, we devised an object model to represent molecular entities, their associated measurements, and the context and provenance relevant to these measurements. KinoML ^13^ is a modular and extensible framework for ML in small molecule drug discovery, with a special focus on kinases. KinoML enables users to easily:

- Access and download data: from online data sources, such as ChEMBL or PubChem, as well as from their own files, with a focus on data availability and immutability. Represented by step 1 in Figure 4.
- Featurize data: so that it is ML readable. KinoML offers a wide variety of featurization schemes (molecular representations that are ML-readable), from ligand-only to ligand:kinase complexes. Represented by step 2 in Figure 4.
- Run structure-based and structure-free experiments: using KinoML’s workflows and ML architectures, a model can be easily trained and tested. These workflows have a special focus on reproducibility. Represented by step 3 in Figure 4.

Currently, the main purpose of KinoML is to help users curate and process kinase data so that it can be used to conduct ML kinase:ligand binding affinity prediction experiments. Tutorials on how to use KinoML, as well as working examples showcasing how to perform experiments end-to-end can be found in the KinoML GitHub repository ^14^. Note that despite KinoML’s focus on kinases, it can be applied to any protein system. For more detailed usage instructions, please refer to the Documentation (https://openkinome.org/kinoml/). Figure 4 displays a typical workflow using KinoML. First, the data is ingested using `DatasetProvider`, which provides a list of `Measurements`. Each measurement contains: the activity values of the measurement, information about the assay conditions of the experiment `(AssayConditions)`, and other `Metadata` for reproducibility purposes. Also, each measurement is associated with a `System`, which contains the information about the specific `Protein` and `Ligand`. The protein and ligand information can be featurized (molecular representation), in such a way that their molecular representations and their associated values can be used for ML applications. Furthermore, KinoML has the `kinoml.ml` module which allows users to easily train and evaluate several integrated ML models.

KinoML is able to achieve the workflow presented in Figure 4 thanks to its object-oriented modular design. The following section provides a more detailed explanation of the object model used in KinoML.

**Figure 4.**
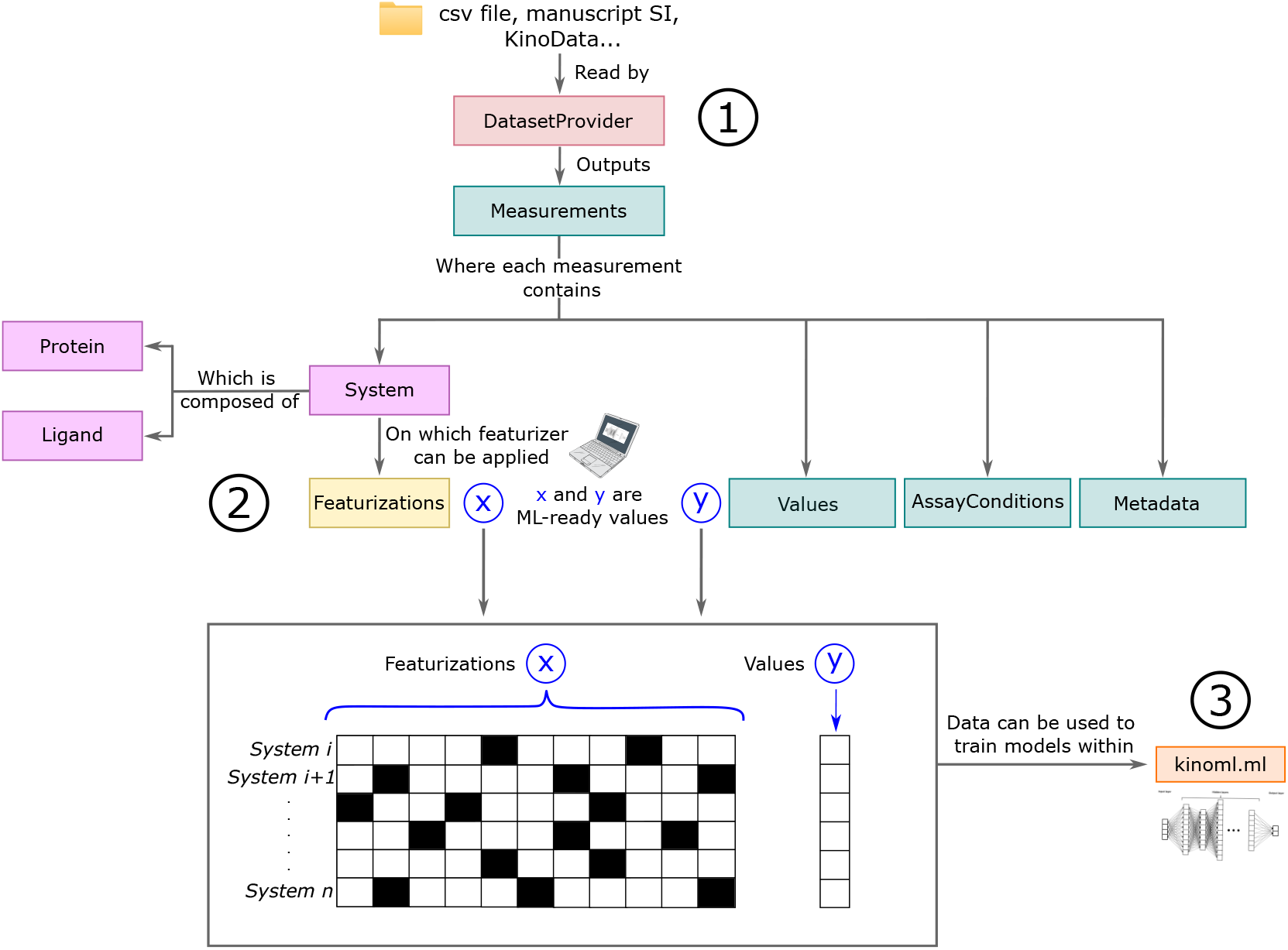
KinoML workflow from data acquisition to featurization. The data is ingested with `DatasetProvider` (step 1), which provides a list of `Measurements`, where each has a `System` and a list of `Values` associated with it. The System information can then be featurized (step 2). Finally, it is possible to use KinoML to easily run structure-based experiments with KinoML’s implemented ML models (step 3). Colors represent objects of the same class.

### KinoML’s object model

KinoML allows users to easily transform molecular systems and their measurements into an ML-compatible format, typically numerically through single or multi-dimensional tensors. To achieve this, the first step is to funnel the clean and processed data into KinoML. This is achieved via the `DatasetProvider` class, depicted in red in Figure 4. `DatasetProvider` can filter and extract data from ChEMBL and PKIS2 csv-files. An example of how a ChEMBL csv file should look like can be found here. This file is also the default kinase ChEMBL csv file in KinoML, so if no url or csv file is provided, then it will use this file obtained from ChEMBL v33.

`DatasetProvider` then reads the ChEMBL or PKIS2 csv-files, processes the data based on UniprotIDs, and extracts their corresponding pIC50, pK__i__, pK__d__, and percentage displacement measurement values. If the user does not want to use the whole dataset, they can specify the UniprotID of the protein(s) of interest, as well as the type of measurement(s) they are interested in (“pIC50”, “pKi” and/or “pKd”), and well as the sample size in case they do not want to use all entries of the dataset.

Once the data is passed through `DatasetProvider`, the data is converted into a list of `Measurements`, which is colored in green in Figure 4. Each `Measurement` has an associated array of values. These values are the different bioactivity measurements that the user wants to work with. The bioactivity measurements that KinoML can work with are: *pIC*50, *pK*__;__, *pK*__*d*__ . Therefore, the array of values will be part of the numbers that the ML model will work with, specifically they will be the y values in Figure 4. Each measurement also has `AssayConditions` and `Metadata` objects associated with it. The `AssayConditions` contains biochemical information regarding the experimental assay from which the measurement was obtained, such as pH or concentration values. The `Metadata` contains information for data provenance and reproducibility, such as the year the measurement was taken or the measurement units.

Each `Measurement` is also associated with a `System` (purple box in Figure 4), which specifies the actual protein and ligands, studied to achieve this measurement. Therefore, the types of molecular components that KinoML can handle are:

- `Ligand`:Ligands (small molecules with certain activity for a target) are encoded as objects within Ki-noML thanks to the `Ligand class`. The `Ligand` is based on the OpenFF-Toolkit [41] `Molecule` object, which can be accessed via the molecule attribute. This allows usage of OpenFF-Toolkit Molecule methods, including conversion to other toolkits, e.g. RDKit [42] and OpenEye (https://www.eyesopen.com/). `Ligands` can be directly initialized via SMILES.
- `Protein`:KinoML provides two types of protein objects: Protein (applicable to all proteins) and KLIFS-Kinase (which allows access to information from the kinase-specific KLIFS database). Similar to Ligand, proteins can be directly or lazily initialized. Again, the molecular structure is accessible via the Molecule attribute. Both protein objects support two toolkits: MDAnalysis (https://www.mdanalysis.org/) and OpenEye (https://www.eyesopen.com/), which can be specified via the toolkit argument. Another important attribute of proteins is their sequence. Note that depending on the desired featurizer, a molecular structure may not be required. For example, in the case of OneHotEncoding only the sequence is necessary. Hence, one can also initialize `Protein` and `KLIFSKi`nase using sequence identifiers only, e.g. UniProt ID or NCBI ID.

Both, `Ligand` and `Protein`, are encapsulated into a `System`, as depicted in Figure 4. This allows KinoML to store all molecular components for a given array of activity `values`. The `System` object can contain just a `Ligand` in case of a purely ligand-based model or featurization, but it can also contain a `Protein`. Therefore, KinoML has implemented three types of systems: `LigandSystem, ProteinSystem`, and `ProteinLigandComplex`.

#### KinoML Featurizers

KinoML provides convenient functions for transforming molecular data into tensors for training models in ML applications. The class `kinoml`.features has several featurizers implemented, for ligands, proteins, and protein-ligand complexes. Examples of these featurizers are: `MorganFingerprintFeaturizer`, `OneHotSMILES-Featurizer` or `OEProteinStructureFeaturizer`. In a nutshell, the `MorganFingerprintFeaturizer` encodes the atom groups of the ligand into a binary vector with length and radius as its two parameters. One hot encoding consists of transforming an array into 0 and 1s, so the `OneHotSMILESFeaturizer` converts the SMILE string representation of a molecule into a one-hot encoding. Finally, the `OEProteinStructureFeaturizer` uses the OpenEye toolkit to prepare a protein structure (modeling missing loops, building missing side chains, etc.) to make the structure ready for docking or to run simulations. A comprehensive explanation of these different featurization schemes can be found in the teaching platform TeachOpenCADD [16]. Furthermore, KinoML has “featurization pipelines”, which are containers for multiple featurizers. This architecture allows for running several featurizations (which depend on each other) in a pipeline to retain as much information from the protein:ligand complex as possible. E.g. given a featurization pipeline where featurization *A* is followed by featurization *B*, this would mean that feature *X* from featurizer *A* can be concatenated with feature *Y* from featurizer *B* to form the resulting feature *XY* . Figure 5 shows a schematic representation of a featurization pipeline applied to a set of kinases and ligands. In this figure, the kinase ABL1 and ligand erlotinib are highlighted to exemplify how the featurization pipeline works in KinoML. In this case, the kinase and ligand are featurized (*x* value) and they are associated with their corresponding affinity (e.g. *pIC*50) value (*y* value). These *x* and *y* values (in combinations with others) can then be used to train an ML model. However, it is important to note that although merging different featurizations into pipelines is essential for the modularity of the framework, this also leads to an increased memory consumption when permanently storing all features generated by the different featurizers of the pipeline. Hence, an option to only store the last featurization for each system is introduced. Jupyter Notebooks showcasing the different featurizers are available within KinoML and a guide on how to use each of them can be found at https://github.com/openkinome/kinoml/tree/master/tutorials/getting_started.

**Figure 5.**
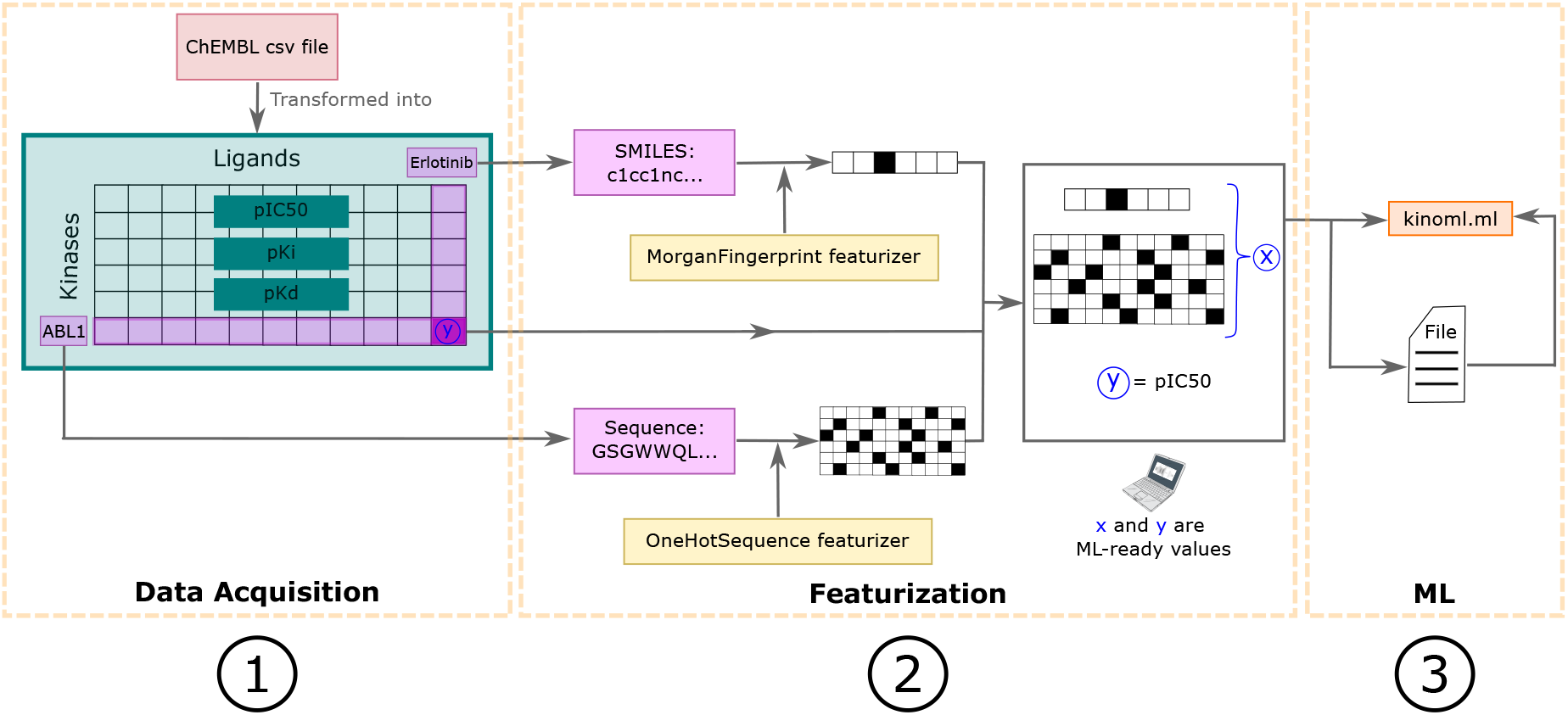
Scalable and customizable pipelines allow featurization of ligand and protein information in a modular and extensible fashion. This is a schematic example of a KinoML workflow from data acquisition to its use in an ML framework. It is the same workflow as depicted in Figure 4. First, in step 1) a ChEMBL csv flle is read by DatasetProvider, which converts the csv file into a dataframe with all ligand-kinase pairs with their respective binding aflnity values of interest. In this case, the values of pIC50, pK_i_, and pK_d_ were stored in the dataframe for each pair. Here, we will only focus on the kinase-ligand pair highlighted: ABL1 and Erlotinib respectively. Proceeding to step 2), the kinase and ligand are featurized with different featurizers. The featurizer transforms the information of the system’s components into numerical arrays, *x*. Here, the ligand and protein components are transformed into fingerprint and one-hot-encoded sequence representations, respectively. Similarly, the binding affinity value associated with this kinase-ligand pair is represented as “y”, and it is the corresponding affinity value for “x”. Finally, as depicted in step 3) these “x” and “y” values are ready to be used for an ML model, and they can be exported to the desired ML framework and saved to disk for later usage. Note that several featurizers already have been implemented providing diverse encoding possibilities for ligands, proteins, and protein:ligand complexes. The modular design of KinoML allows for straightforward design of new featurization pipelines.

### Structural representations

KinoML allows the user to easily apply docking pipelines to protein:ligand systems. KinoML has interfaces to several docking and template docking methods, that allow users to prepare protein structures and to dock small molecules into their binding sites. The docking strategies currently covered within KinoML use OpenEye [43] or Schrödinger software (https://newsite.schrodinger.com/). ^15^ Structural featurizers can output the resulting protein in PDB format and a prepared and/or docked small molecule in SDF format.

#### Docking algorithms in KinoML

Docking is a computational molecular modeling technique used to predict the preferred binding orientation or pose of a small molecule (ligand) within a binding site on a target protein. [44] Docking is classically a two-step process of positioning and scoring the ligand within the binding site to suggest the most energetically favorable binding mode. This can provide valuable insights into the structure activity relationship and key protein:ligand interactions. However, issues such as inaccurate scoring functions and the flexibility of both ligands and proteins can limit the accuracy of docking predictions. Template docking, which uses known ligand-protein complexes to guide the docking process, can help address some of these limitations by providing a more reliable framework for predicting binding poses.

For systems with one protein and one ligand, the docking pipelines implemented in KinoML are:

- OpenEye FRED (Fast Rigid Exhaustive Docking): this docking method rapidly explores ligand conformations and orientations within the binding site. This is known as a standard docking protocol.
- OpenEye Hybrid: this docking method can use either a single receptor structure or multiple structures of the target protein. For single structure docking, it requires the receptor and ligand database. For multiple structures, it requires all receptor files and the ligand database. Hybrid focuses on leveraging multiple protein structures to improve docking accuracy.
- OpenEye POSIT: this pose-prediction tool assumes similar ligands bind similarly [45]. It selects the best docking method for a ligand and estimates the probability that the pose is within 2.0 Å of the actual pose. POSIT uses known bound ligands to guide docking and optimize geometry, focusing on the accuracy of the predicted binding pose.
- Schrödinger Glide [46]: employs hierarchical filters and grid representations for ligand pose scoring.

Listing 1 illustrates how to apply KinoML’s featurization Pipeline to perform docking on a ligand:kinase system. In this example, the protein is loaded by specifying the UniprotID and the ligand is initialised via SMILES. Then, the most suitable PDB structure (ligand binding mode) to dock into is found with `MostSimilar-PDBLigandFeaturizer`. Template docking is a popular docking technique that centers around utilizing the pose information of the template ligand as a guide for predicting the binding mode of the target ligand. For this purpose, `MostSimilarPDBLigandFeaturizer` is implemented within KinoML, which finds the most suitable structure for docking in the PDB based on ligand similarity. This is particularly useful for larger sets of ligands, for which manually specifying the most suitable PDB structure to dock into would be very time consuming. `MostSimilarPDBLigandFeaturizer` allows the user to choose from several similarity metrics: fingerprints, most common substructure, and OpenEye’s and Schrödinger’s shapes. Then, as exemplified in Listing 1, once the most suitable structure to dock into has been found, the docking on this pose is run with `OEDockingFeaturizer`. Therefore, with KinoML a molecular system can be initialized and featurized in just a few lines of code.

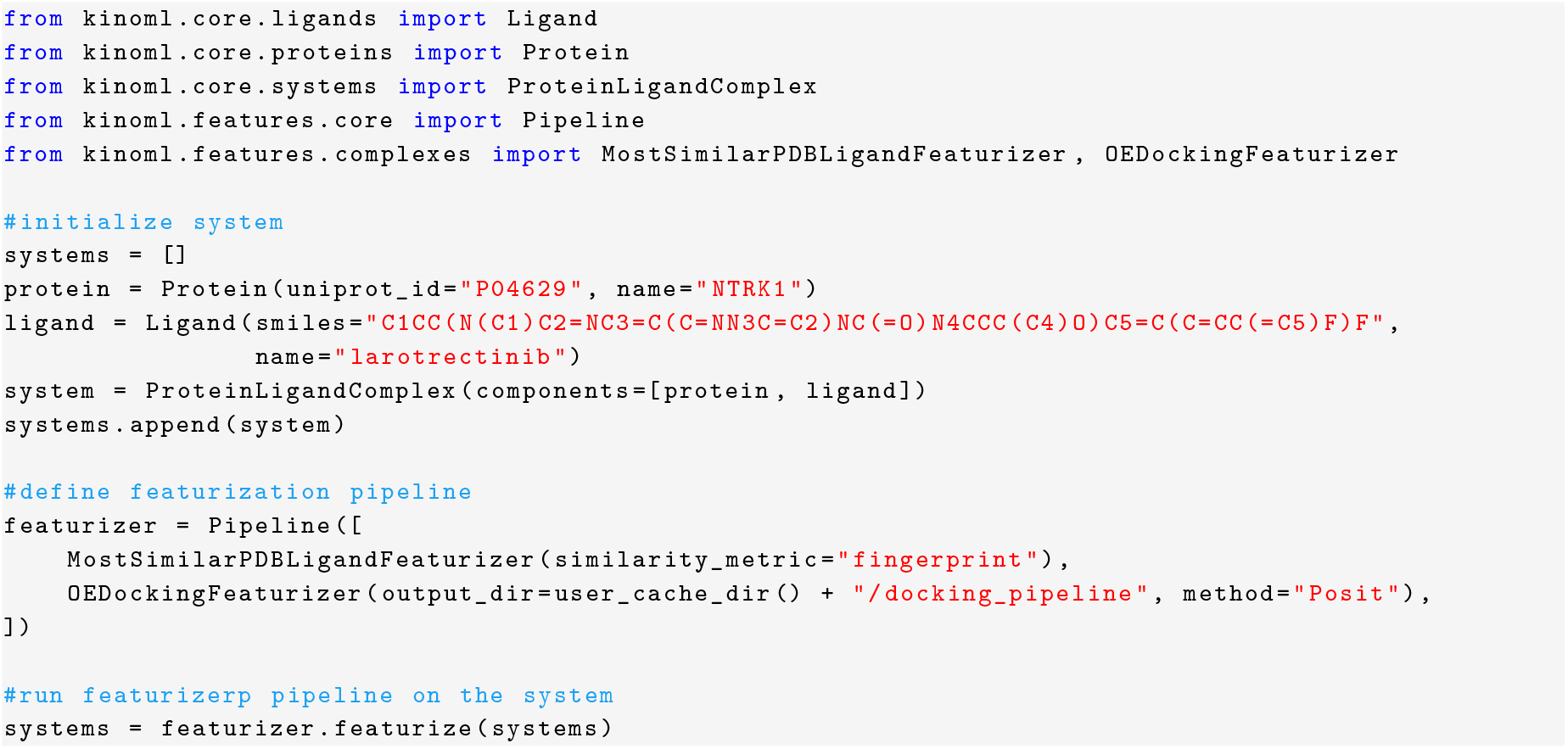

**Listing 1**. Code snippet showing how to use the KinoML featurization pipeline to apply several featurizers on a system.

For a more detailed explanation of all the featurizers implemented within KinoML and a guide on how to use each of them, please refer to the following tutorial folder: https://github.com/openkinome/kinoml/tree/master/tutorials. In this folder you will find several notebooks showcasing KinoML featurization capabilities. Overall, KinoML’s object model allows users to easily fetch data from ChEMBL or PKIS2 data sources and then to featurize this data so that it can be used for ML purposes.

### Exposing tensors to machine learning frameworks

The featurization of the molecular components are represented as tensors. The primary aim of storing these molecular feature tensors is to ensure they can be easily accessed and reused for training and evaluating machine learning models. Archiving these tensors facilitates reproducibility and allows for consistent benchmarking across different studies.

Machine learning feature tensors should be kept on disk for archival, reusability, and reproducibility of the trained networks. These tensors can be stored in a platform-agnostic way as long as the corresponding adapters to popular machine learning frameworks are in place. The chosen output format should offer high-performance read and write access, without hindering the flexibility to support heterogeneously shaped tensors. Given the expected tabular nature of the data and its diversity in the KinoML pipelines, Parquet files (https://parquet.apache.org/) were used to store the feature tensors, which have been proven to be a efficient and flexible format, allowing nested column data and compression techniques specific to each column. The Parquet format is also a free and open source, making it a good candidate for portability due to its support in different programming languages and machine learning frameworks.

Our first approach was to serialize the featurized tensors to a Numpy array. However, Numpy arrays cannot be used for data with heterogeneous shapes.

Therefore, we dropped the Numpy-only design requirement and chose another data structure with better support for disk serialization. The awkward-array project provides a Pythonic interface to ragged arrays (tensors composed of arrays of varying dimensions) with acceptable performance for our domain. By using awkward-arrays, we could leverage Parquet files for storage. Parquet’s compatibility with awkward-arrays addressed the limitations of our earlier attempts by enabling efficient serialization of ragged arrays. Moreover, Parquet’s flexibility allowed us to store arbitrary metadata next to each data set entry, providing extensibility for future needs.

Using ragged arrays allowed us to address the challenge posed by heterogeneous tensor shapes, and they are a good balance between performance and flexibility. This design choice ensures efficient read and write access while supporting diverse featurization schemes. However, this approach does add an extra layer of complexity into the data handling and an increased storage cost.

### Ensuring reproducible molecular machine learning

Throughout the different steps of this data journey to build KinoML, we tried our best to ensure every action could be reproduced at a later point in the future. This section will consolidate each of these “checkpoints” to ensure reproducibility that are scattered across the manuscript into a list of key ideas.

- **Re-executable data manipulation**:ensure that every programmatic manipulation of the data is re-executable. Always eliminate machine-dependent states and isolate and freeze execution environments, including by documenting the dependencies and versions of libraries. It is also important to use relative paths and to document options and non-obvious code blocks. Furthermore, ensure determinism by fixing random seeds across all libraries and runtimes, this step guarantees that random processes, such as data splitting or model initialization, remain consistent across different executions. Lastly, make sure to test in other computers or, even better, use automated pipelines to ensure reproducibility over time.
- **Ensure consistency in model building and initialization**:Ensure that models are being built in the same way, to ensure consistent reproducible results. In order for users to build featurization pipelines and train models with KinoML, they will just need to modify a hyperparameter file where all featurization and model details will be specified. This way, users can quickly compare if parameters are consistent across models.
- **Ensure consistent and traceable training operations**:It is important to ensure that splitting, batching, and random seeds are consistent during benchmarking and setup. For non-random algorithms, ensure deterministic results to maintain model consistency. In KinoML, this will be specified in a hyperparameter file so users can easily check the parameters they have inputted and ensure they are consistent across runs.
- **Utilize MLOps platforms to track development progress and ensure all training and results are recorded**:MLOps platforms are tools and frameworks designed to manage the end-to-end machine learning lifecycle, including development, deployment, monitoring, and governance. Utilizing these platforms helps ensure that there is a record of all training and results, key for consistency checks and reproducibility. Examples of MLOPs platforms are TensorBoard ^16^ or Weights & Biases ^17^.
- **Standardize evaluation protocols for ML models**:To ensure valid comparisons and observable improvement in model construction, it will be important to generate a standard pipeline for evaluating model results. This includes having a consistent metric (e.g., accuracy, precision, recall, F1-score, AUC-ROC) for evaluating all models, uniform data splits for evaluating different models or to standardize the process of analyzing model errors and misclassifications.

### KinoML example of end-to-end usage

KinoML can already be used to run binding affinity experiments, from data curation to affinity prediction using one of KinoML models. In the KinoML documentation we provide examples of how to run experiments end-to-end using KinoML. Detailed examples of this can be found here. KinoML is build in such a way that the user only needs to change the parameters of two hyperparameter files, one contains all the featurization pipeline information, and the other one contains the model training information.

The featurization hyperparameters file contains information such as the dataset provider, the measurement types of interest, the sample size or the featurizers that the user wants to apply on the data. On the other hand, the model training hyperparameters file contains fields such as the model name, number of epochs or batch size. This way, users only need to modify these files accordingly, and follow the same workflow as specified in the tutorials to run any experiment on interest.

Furthermore, KinoML has already been used in publications. For example, Backenköhler et al. [9] and Schaller et at. [13] have both used KinoML to create featurization and docking pipelines of their kinase data, which were then used for ML applications. KinoML also served as the basis for some work in the Chodera lab within the Covid Moonshot context [47], and it inspired the software development in the ASAP Discovery project^18^. Additionally, KinoML’s generalized framework has been utilized to model conformations of PDB structures [48]. Moreover, KinoML provided fingerprinting frameworks for classifying inhibitors, which led to the identification of combinations of kinase inhibitors that exhibit reduced off-target efficacy [49].

## Conclusions/Future Perspectives

In this work we have presented our experiences building KinoML, highlighting its capabilities, usability, and its focus on ensuring reproducibility in ML experiments. However, the purpose of this manuscript is not only to display KinoML’s potential, but also to establish some best practices when conducting molecular ML experiments. The summary of the key recommended procedures discussed in this manuscript are:

- **Reproducible data collection**:to always ensure reproducibility and immutability of the starting point of your experiments, store unmodified raw data and used versioned database copies.
- **Data deposition**:for any data preprocessing step before storing it (e.g. extracting data from HTML), always ensure to provide the code used for this purpose. Also, choose where to store your data depending on its size keeping in mind to guarantee long-term availability of the stored data.
- **Molecular data collection**:when collecting and searching for protein data across different data sources it is best to use the *UniprotID* as the primary search key, since this is universal. In the case of kinases, we have shown that different kinase-centric databases have different kinase definitions and names, so it is important to craft a unified database with the target kinases of interest. Another main aspect when collecting molecular data is to ensure the harmonization of the bioactivity data. Pay special attention to the units and assay conditions of all measurements to ensure that either they can be converted to or have the same units for appropriate comparison.
- **Data format for ML:**tensors representing the molecular featurizations should always be kept on disk for reusability and reproducibility of ML models. It is important to choose a tensor output format that can support heterogeneously shaped tensors. For example, in KinoML we chose ragged arrays, which can store heterogeneous tensors and offer a good balance between performance and flexibility.

By implementing the best practices discussed above, KinoML presents itself as a machine learning framework for small molecule drug discovery with a strong focus on reproducibility.

Overall, KinoML serves as a comprehensive and modular ML framework that not only facilitates easy access and curation of data but also enables the featurization of data, making it ML-readable in a user-friendly fashion. By seamlessly integrating structure-based experiments, KinoML has the potential to be a powerful tool for researchers in the field of kinase drug discovery. For example, KinoML’s docking pipelines have been used in published work. [9, 13]. Future work regarding KinoML is the incorporation of physical methods, such as scalable free energy calculations, for more precise and physically-informed workflows for engineering small molecule kinase inhibitors with specific polypharmacologies. Note that KinoML is under heavy development right now, and we hope an official release is forthcoming.

The lessons learned from KinoML’s construction are applicable beyond the kinase superfamily, offering insights and guidance for the development of infrastructure in other areas of drug design. Furthermore, KinoML is an example of taking steps towards integrating structure-based methods in drug discovery pipelines. Overall, KinoML can serve as a key tool, facilitating and guiding users toward more efficient, reproducible, and impactful research in the realm of structure-enabled ML drug design.

## Code and data availability

All code is available under the MIT license at the openkinome organization https://github.com/openkinome. The acquisition is available under https://github.com/openkinome/kinodata, the remaining functionality under https://github.com/openkinome/kinoml.

## Author Contributions

## Acknowledgments

The authors thank OpenEye Scientific Cadence Molecular Sciences for providing a free academic license to the OpenEye Toolkits to the Chodera lab.

## Footnotes

1 https://www.discoverx.com/

2 https://lincs.hms.harvard.edu/kinomescan/

3 https://www.eurofinsdiscovery.com/solution/kdelect

4 https://www.ebi.ac.uk/chembl/

5 https://pubchem.ncbi.nlm.nih.gov/

6 https://github.com/openkinome/kinodata

7 https://www.ebi.ac.uk/chembl/

8 https://pubchem.ncbi.nlm.nih.gov/

9 https://github.com/openkinome/kinodata/blob/master/kinases-in-chembl/kinases_in_chembl.ipynb

10 https://zenodo.org/

11 https://figshare.com/

12 https://github.com/openkinome/kinodata/blob/master/kinase-bioactivities-in-chembl/kinase-bioactivities-in-chembl.ipynb

13 https://github.com/openkinome/kinoml

14 https://github.com/openkinome/kinoml/tree/master/tutorials

15 Note, that a valid license for at least one of the two is needed to use KinoML docking capabilities. As of September 10, 2024, OpenEye offers to apply for free academic licenses.

16 https://www.tensorflow.org/tensorboard

17 https://wandb.ai/site

18 https://github.com/asapdiscovery.

